# Implementing a framework of carbon and nitrogen feedback responses into a plant resource allocation model

**DOI:** 10.1101/2023.06.27.546727

**Authors:** Bethany L Holland, Nicholas A M Monk, Richard H Clayton, Colin P Osborne

## Abstract

- The allocation of resources to roots and shoots can greatly alter total plant mass. Allocation is thought to be the consequence of growth processes (i.e uptake rates, transport rates, growth rates) and the communication between them via signalling mechanisms. Feedbacks that alter growth processes are induced in nature by changes in the internal pools of carbon and nitrogen, but how these function together to define allocation remains unclear.
- We introduce a framework model of internal feedback responses to changes in plant carbon and nitrogen concentrations. We evaluate how well the model responds to changes in carbon and nitrogen availability by simulating external environmental perturbations that influence the uptake of resources.
- The model reflects experimental results when looking at the effect of atmospheric *CO*_2_ and soil nitrogen concentrations on total plant mass and replicates observed responses to leaf defoliation events. Overall this shows that a combination of known signalling mechanisms are sufficient to reproduce experimentally observed responses to external resource availability.
- Model simulations highlight key areas of uncertainty where more empirical data is needed. In particular, quantitative data is needed to establish the strengths and rates at which feedback responses to carbon and nitrogen substrate concentrations alter growth and uptake rates.

## 1. Introduction

Changes to the allocation of resources to different plant tissues (e.g. leaves, roots, stem and seeds) greatly impact total biomass and crop yield and arise when plants react to changes in the environment. The responses of biomass allocation to the environment are thought to balance the uptake of carbon and nitrogen. Crop yields depend on the coordinated acquisition of carbon and nitrogen by the leaves and roots respectively and the coordinated use of these resources within each part of the plant. Carbon and nitrogen assimilation and the use of their products are entirely interdependent (Moorby, 1977; Paul and Foyer, 2001; Kaschuk et al., 2010). For example, the energy required for nitrogen assimilation is provided via photosynthesis (Stitt et al., 2002) and byproducts of nitrogen assimilation are required for photosynthesis to occur (Zhu et al., 2008). Gaining a quantitative understanding of how carbon and nitrogen behave together in plant metabolism and signalling can therefore elucidate how such responses can be optimised to enhance plant growth.

Environmental variation alters processes within the plant required for growth (respiration, photosynthesis, nutrient uptake, etc.) to differing extents, meaning that different internal processes become more or less limiting under different environmental conditions. This imbalance necessitates a balancing of energy producing and utilising processes which is modulated by molecular regulation (Paul and Foyer, 2001). In particular, intermediate products from carbon and nitrogen assimilation such as nitrate, sugars, and amino acids reflect the carbon:nitrogen status of the plant and act as signals (or feedbacks) for gene expression to affect many cellular processes (Figure 1). This leads to important interactions between the signalling pathways for carbon and nitrogen. However, thousands of genes respond to changes in sugar concentrations (Lastdrager et al., 2014). A simplification of these processes is therefore needed to understand how they interact at a whole plant scale.

**Figure 1:**
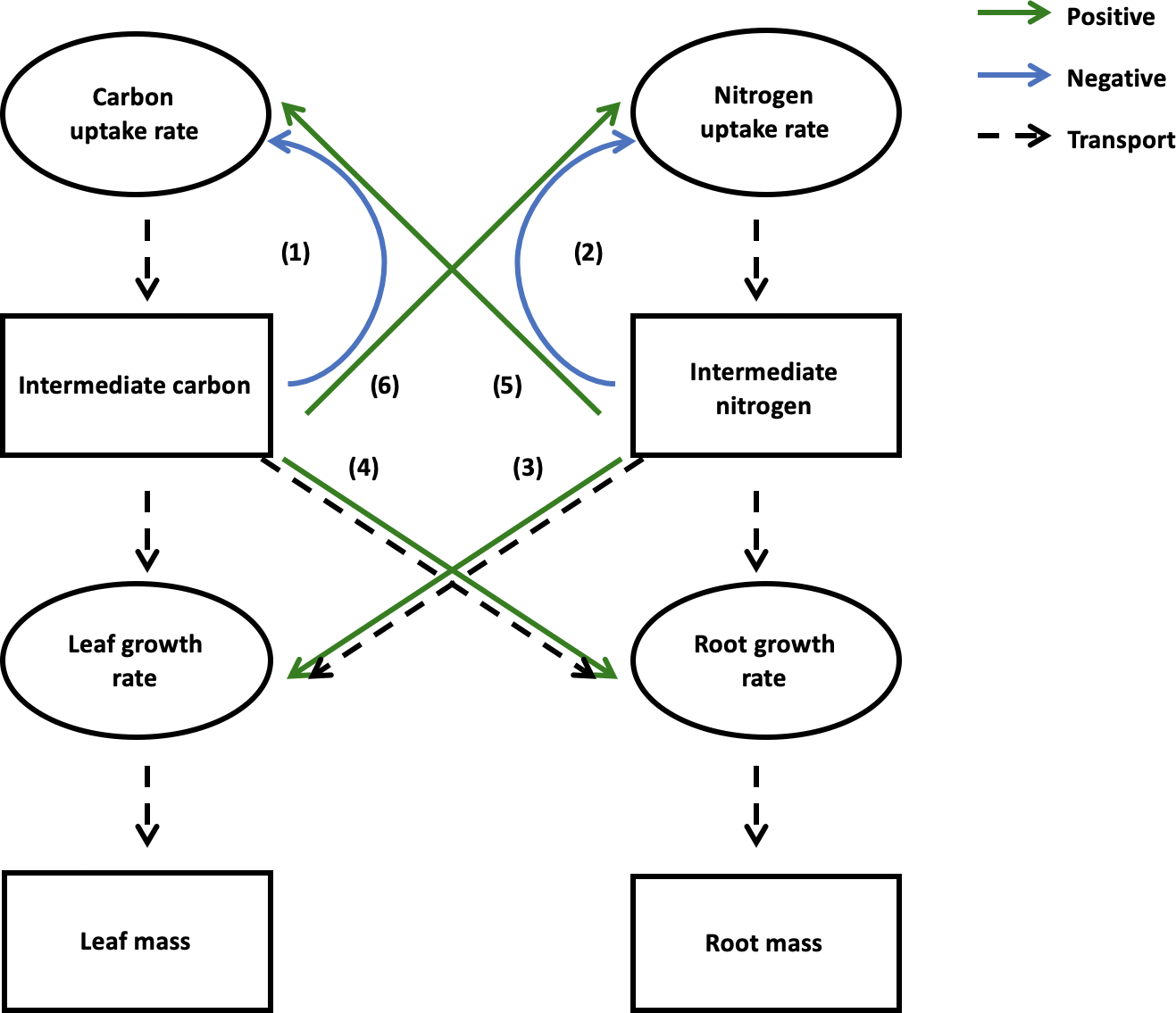
Summary of feedback responses to carbon and nitrogen concentrations observed experimentally. Rectangles represent products of growth processes (e.g. pools of C and N and leaf and root mass) and ovals represent the internal processes of resource uptake and growth. Dashed lines represent the transport of resources between compartments. Blue arrows are negative feedbacks and green are positive feedbacks. Evidence for each of these is reviewed in the text. 1. High leaf carbon concentrations decrease carbon uptake rate. 2. High root nitrogen concentrations decrease nitrogen uptake rate. 3. High leaf nitrogen concentrations increases leaf growth rate. 4. High root carbon concentrations increase root growth rate. 5. High leaf nitrogen concentrations increase carbon uptake rate. 6. High root carbon concentrations increase nitrogen uptake rate.

### Carbon assimilation

The rate of photosynthesis is sensitive to leaf carbon concentrations. Increasing atmo-spheric *CO*_2_ typically enables the build up of starch and other carbohydrates in the leaves in a matter of hours. Via a number of mechanisms that sense leaf carbohydrate status, this triggers an immediate reduction in rubisco activity which is an important constraint on carbon assimilation (Paul and Foyer, 2001). There is debate as to whether high starch concentrations weaken chloroplast function or not (Paul and Foyer, 2001). However, a large body of evidence (Paul and Foyer, 2001; Smith and Stitt, 2007; Kelly et al., 2013; White et al., 2016) shows that high carbon concentrations have a negative feedback on carbon uptake (Feedback 1, see Figure 1). For example sucrose has been shown to reduce the transcription of photosynthesis related genes (Graham, 1996; Koch, 1996; Chan and Yu, 1998; Sheen et al., 1999; Smith and Stitt, 2007). Excess sucrose is sensed by hexokinase which also triggers the closure of stomata, leading to a reduction in photosynthesis (Kelly et al., 2013).

Cytokinins in the root are very sensitive to nitrogen supply and the transport of this hormone from roots to leaves promotes the expression of genes linked to photosynthesis. This implies that high nitrogen status in the roots promotes an increase in photosynthesis (Feedback 5, see Figure 1) (Paul and Foyer, 2001).

### Nitrogen uptake & assimilation

In nitrogen assimilation, nitrate is often converted into ammonium via nitrate reductase (NR) in the cytosol and the chloroplast in the leaves but this can also occur in the roots of some plant species (Weissman, 1972). NR is synthesised in the presence of nitrate and is degraded when it is absent. When nitrogen supply to the roots is limited, the synthesis of ammonium and consequently glutamine and glutamate synthesis (via the glutamine synthetase (GS) and glutamate synthase (GOGAT) pathway (Lam et al., 1996)) occurs in the plastids of the roots (Raven et al., 1999). Nitrogen assimilation is carbon-dependent because energy is required for the synthesis of glutamate and glutamine and is a key stage where carbon metabolism and nitrogen metabolism interact (Hodges, 2002).

Plant sugars can be used in place of light to induce the genes responsible for NR in the leaves (Cheng et al., 1992; Klein et al., 2000; Iglesias-Bartolomé et al., 2004; Reda, 2015) consequently increasing nitrogen uptake rate. Reda (2015) shows increased NR activity following sugar treatments of 8 hours but there is no further evidence about how this mechanism works. The location where nitrogen derived signals are sensed alters the type of feedback on nitrogen uptake rate. Not only do sugars and inorganic acids stimulate multiple stages of nitrogen assimilation but products of nitrogen assimilation (glutamine and glutamate) act as signals for the expression of genes responsible for the inhibition of NR and therefore reduce nitrate uptake in the leaves (Siddiqi et al., 1990; Clarkson and Lüttge, 1991; Muller and Touraine, 1992; King et al., 1993; Rufty et al., 1993; Imsande and Touraine, 1994; Reda, 2015). In the roots, glutamine and glutamate induce NR activity (Shaner and Boyer, 1976; Wray, 1993; Gojon et al., 1998; Reda, 2015). Nitrate induces genes responsible for NR within 30 minutes but this is only when photosynthesis is active.

In summary, experimental evidence indicates two feedbacks: firstly, the increase in nitrogen uptake and assimilation when internal carbon concentration is high (Feedback 6, see Figure 1) and secondly, the reduction in nitrogen uptake when nitrogen concentration is high (Feedback 2, see Figure 1).

### Growth rate

When carbon assimilate availability is high, new sinks can be formed (Paul and Foyer, 2001). It has been found that sucrose is important in the regulation of plant growth such that it induces the production and transport of auxin (hormone responsible for growth), therefore increasing sink activity (Lilley et al., 2012; Sairanen et al., 2012; Stokes et al., 2013). Xiong et al. (2013) found that root glucose activates TOR protein kinase, promoting the activity of root meristems. This represents a positive feedback on root growth when the concentrations of carbon intermediates are high (Feedback 4, see Figure 1).

The allocation of carbon within the plant is affected by nitrogen availability (Nunes-Nesi et al., 2010), such that high plant nitrogen content increases shoot:root growth (Stitt and Krapp, 1999). Scheible et al. (1997) show that the presence of nitrogen in the roots increases protein synthesis and root growth rate but shoot growth rate is higher, leading to a stronger allocation of growth towards the leaves. This identifies a further feedback to increase leaf growth when nitrogen concentrations are high (Feedback 3, see Figure 1).

### Modelling feedbacks

Many models simulate growth by considering the dependence of growth on carbon and nitrogen supply (i.e. growth is substrate limited) but do not necessarily include the signalling feedbacks on uptake and use (Thornley, 1972; Hunt et al., 1998; Bartelink, 1998; ÅAgren et al., 2012; Cheeseman, 1993; Siddiqi and Glass, 1986; Shaw and Cheung, 2018). The models which do simulate resource dependencies of uptake rates or growth are limited by either not considering above and below ground material (ÅAgren et al., 2012), using a functional balance assumption (Bartelink, 1998; Hunt et al., 1998; Shaw and Cheung, 2018) or only focusing on leaf canopies or root systems (Dunbabin et al., 2002; Pao et al., 2018). Carbon- and nitrogen-derived signals have been observed in plant growth, but how these signals are coordinated together with changes in nutrient availability is still unknown. This raises questions such as: 1. How fast is the feedback? 2. Is this a feedback that is turned on or off or does it happen incrementally? 3. If it does work like a switch, what threshold values cause this feedback? 4. Can all of the known feedbacks operate simultaneously to generate a stable system?

In many cases, part of the mechanism has been elucidated, but the complete sequence of events needed to simulate the genetic interactions is unknown. Even when mechanisms are known, further data would be needed to parameterise model functions. Most published work on the mechanisms compare gene expression and / or changes in enzyme activity for a plant with and without sugar or amino acid treatments. Some acknowledge the time it takes for a gene to be expressed (within 30 minutes (Reda, 2015)) or the time taken for resources to accumulate (Paul and Foyer, 2001) and most acknowledge a time length of the treatment (Reda, 2015). Some work does provide a timescale. For example, Xiong et al. (2013) show that meristem activation occurs within 24 hours of treating seedlings with glucose. However, there is little information on how long it takes from the accumulation or depletion of carbon or nitrogen and the expression of genes, to the increased enzyme activity which represents the full feedback process. Without this information, it is not possible to compare the rates at which the various feedbacks operate in a parameterised model.

Though some previous models have simulated the dependency of source activity on carbon and nitrogen concentrations, no previous models have attempted to simultaneously simulate feedbacks on source and sink activity. It is unknown how the unification of multiple feedbacks alters growth allocation individually and collectively. This paper explores the mechanisms responsible for allocation with three objectives. First, to evaluate whether the combination of known signals is sufficient to reproduce empirically observed responses to imbalances in carbon / nitrogen supply. Secondly, to sharpen questions about the unknown aspects of signalling. e.g. rates, thresholds and the nature of signals. Thirdly, to formulate a stable, working model that is capable of reproducing whole-plant signalling and allocation behaviour in response to variable carbon and nitrogen supplies, and that may be adopted for use in crop and vegetation simulation models.

To evaluate the model, two tests have been devised based on empirical observations of the interactive effects of carbon and nitrogen supply on plant functioning and allocation. Experiments show that high soil nitrogen increases the positive effect of atmospheric *CO*_2_ on total plant mass (Coleman et al., 1993; Farage et al., 1998; Kirschbaum and Lambie, 2015), whilst high levels of *CO*_2_ reduce overall plant nitrogen content (Cotrufo et al., 1998; Curtis and Wang, 1998; Norby et al., 1999; Jablonski et al., 2002; Ainsworth and Long, 2005; Taub et al., 2008) and net carbon uptake in relation to intercellular *CO*_2_ (Coleman et al., 1993; Ainsworth et al., 2003). Additionally, reducing carbon source size by defoliation increases leaf nitrogen and the carbon uptake rate of the remaining leaves, whilst reducing intermediate carbon concentrations in the leaves (Rogers et al., 1998).

The aim of this paper is to determine whether the unification of certain carbon & nitrogen feedback responses can reflect whole plant allocation patterns and evaluate the extent to which the parameterised model of feedback responses on growth processes can reproduce these experimentally observed behaviours qualitatively. The model results are compared qualitatively against the results from experimental papers since the model is not calibrated for one specific plant species but instead is parameterised generally for herbaceous plants. This paper shows that the model mostly reacts to changes in *CO*_2_ and nitrogen availability in the same way as experiments carried out on plants, providing a framework to further investigate the dynamics between internal feedback mechanisms underpinning allocation.

## 2. Description

There are typically two main approaches to simulating allocation of resources. The first focuses on reconstructing entire metabolic networks and the flows of energy and materials within them (i.e flux balance analysis models (Shaw and Cheung, 2018; Moreira et al., 2019)) but lacks the feedbacks on metabolism described above, since the details of the molecular mechanisms are not fully known. The second takes a much higher level approach, aiming to reproduce the outcome of the feedbacks in terms of allocation to roots and shoots, and the rates of carbon and nitrogen assimilation, in relation to environmental limitation. This second class of models is phenomenological or teleonomic (Thornley and Johnson, 1990; Buckley and Roberts, 2006; Feller et al., 2015). This approach doesn’t explicitly consider the internal feedbacks that give rise to the behaviour. The model described here operates at an intermediate level. Since the detailed molecular mechanisms are not fully known, we approach this problem by explicitly simulating the behaviour of experimentally observed physiological feedbacks in a qualitative way. The model is thus mechanistic at a physiological but not molecular scale.

For the purposes of the model, plant mass is compartmentalised into two tissue types: leaves and roots, and each tissue type has an intermediate pool of non-structural carbon (i.e. sugars, starch) and nitrogen (i.e. nitrate, ammonium, amino acids). These intermediate products of carbon and nitrogen assimilation are transported around the plant, may accumulate in tissues, and are utilized by tissue growth processes and increase in size when the amounts of carbon or nitrogen taken up into the plant are higher than the amount required for growth.

The model used here is a unification of a widely used and tested transport resistance model (Thornley, 1972) and a new framework of internal feedbacks of internal carbon and nitrogen concentrations on growth processes. The Thornley (1972) model is used since it is an excellent framework to investigate source-sink dynamics and has been used for a variety of different plant species and environmental conditions (Wann and Raper Jr, 1984; Rastetter and Shaver, 1992; Minichin et al., 1994; Dewar et al., 1994).

The model simulates four compartments with two substrates (carbon and nitrogen) moving between the leaf and root, which is represented by a system of 6 first order ordinary differential equations (ODEs) and are available in the supplemental information. The first four equations represent the four pools of substrate (intermediate carbon and nitrogen concentrations in the leaves and roots) and the final two equations represent the masses of leaf and root tissue (Fig. 2). All figures in this paper are produced via MATLAB and the code is available through github (Github repository).

**Figure 2:**
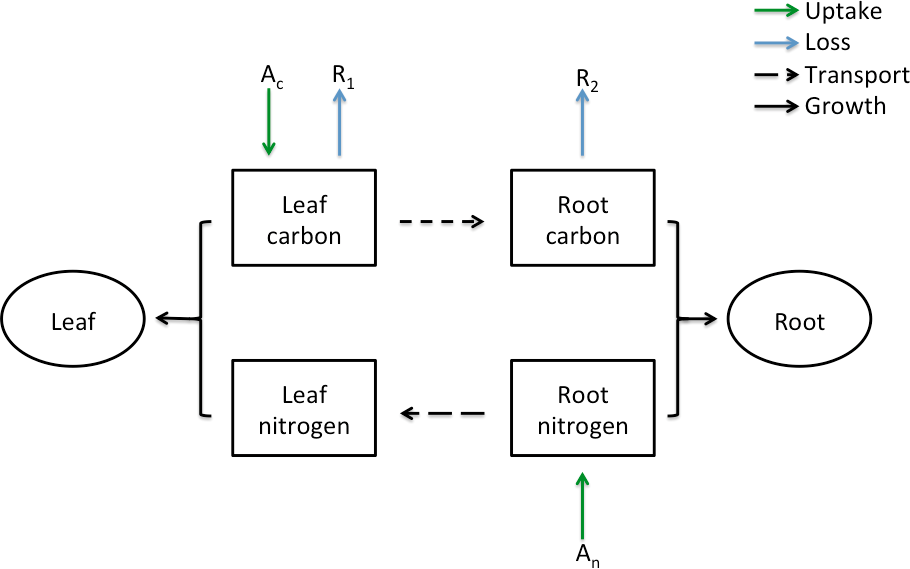
Diagram of the Thornley (1972) model with maintenance respiration. The boxes represent intermediate carbon and nitrogen concentration per unit leaf or root mass and the circles represent total leaf or root mass. Leaf carbon and root nitrogen concentrations increase via carbon (*A_c_*) and nitrogen (*A_n_*) uptake rates per unit leaf or root mass (green arrows). Leaf carbon and root carbon are reduced via leaf (*R*_1_) and root (*R*_2_) maintenance respiration (blue arrows). The black dashed arrows represent transport of resources and the black solid arrows represent the use of resources for growth of leaf and root mass.

The model is based on a combination of the assumptions from Thornley (1972) and additional assumptions to simulate a plant which is sensitive to external and internal fluctua-tions of carbon and nitrogen. First, Thornley assumes that growth of new plant volume is dependent upon the use and transport of substrate.

### Use of substrate & allocation

The rate of use of substrate for growth is derived from bisubstrate enzyme kinetics (Dixon and Webb, 1964 - taken from Thornley (1972)) in which the rate of use of carbon is defined as:

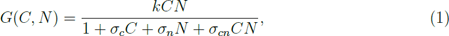

where *k* is the Michaelis-Menten constant (*m*^3^*s^−^*^1^(*kgmol*)*^−^*^1^) and *C* is carbon concentration, *N* is nitrogen concentration. *σ_c_*, *σ_n_*, and *σ_cn_* are rate constants which determine the concentrations at which carbon ((*m^−^*^3^*kgmol*)*^−^*^1^), nitrogen ((*m^−^*^3^*kgmol*)*^−^*^1^) and both carbon and nitrogen ((*m^−^*^3^*kgmol*)*^−^*^2^) start to saturate. This function represents the amount of carbon and / or nitrogen used for the growth of new plant tissue. This allows the growth of leaves and roots to depend upon both carbon and nitrogen, such that allocation of growth to above or below ground biomass is a consequence of changes in intermediate concentrations. For simplicity, the Michaelis-Menten constants for carbon and nitrogen are assumed to be equal (See Eq. (5) & (6)).

### Continuous growth

There is no litter production (i.e. tissue turnover) within the model; the only loss term is maintenance respiration which reduces leaf and root carbon pools. Growth respiration is accounted for by an efficiency constant (*Y_g_*), assuming that a proportion of carbon and nitrogen resources available for growth are lost in order to construct new biomass. The production of leaf and root mass is expressed as exponential growth and therefore does not reach steady state. This represents a stage of vegetative growth which will only stop when relative growth rate (RGR) becomes zero. For this to occur in the model, carbon or nitrogen pool sizes in the leaves must be zero and carbon or nitrogen pool sizes in the roots must be zero.

### Environmental dependence & uptake rates

Thornley (1972) assumes that carbon and nitrogen uptake rates are constant per unit shoot or root volume. Here, this assumption is modified so that carbon uptake rate is also dependent upon atmospheric *CO*_2_ (Eq. (2)) and nitrogen uptake rate upon soil nitrogen concentration (Eq. (3)). This allows the plant to be responsive to environmental events which may cause changes to external carbon and nitrogen concentrations.

For the model to respond to changes in environment, gross carbon uptake rate is modified to become dependent upon atmospheric *CO*_2_. Originally a constant rate (Thornley, 1972), carbon uptake rate (*A_c_*) becomes:

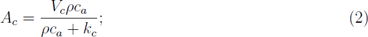

where *V_c_* is maximum carbon uptake (40*µmolm^−^*^2^*s^−^*^1^ (Sage, 1994)), *ρ* is the conversion factor of atmospheric to intercellular carbon (0.7 *µmolmol^−^*^1^*m*^3^ (Katul et al., 2000)), *c_a_*is atmospheric *CO*_2_ (*ppm m^−^*^3^) and *k_c_* is the concentration of *CO*_2_ at half of *V_c_* (200*µmolmol^−^*^1^(Farquhar et al., 1980)).

Nitrogen uptake rate can also be described using the Michaelis-Menten equation (Youngdahl et al., 1982), such that it becomes dependent upon soil nitrogen availability, therefore nitrogen uptake rate (*A_n_*) becomes:

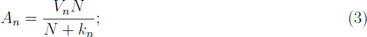

where *V_n_* is maximum nitrogen uptake rate (61*µmolkg^−^*^1^*s^−^*^1^(Youngdahl et al., 1982)), *N* is soil nitrogen content (*µM*) and *k_n_* is soil nitrogen content at half of *V_n_* (103*µM* (Youngdahl et al., 1982)).

### Transport of substrate

The transport of substrate is assumed to follow Münch mass flow such that the amount of intermediate carbon or nitrogen transported between leaves and roots is determined by the difference in their concentrations in source and sink, whilst transport resistance is scaled with plant size (Münch, 1930). As the plant increases in size, the level of transport resistance increases. Thornley (1972) assumes that it takes approximately one day for intermediates to be transported from source to sink. This is reduced to roughly 3 hours in the modified model to increase relative growth rate.

### Internal feedback mechanisms

A framework of internal feedback mechanisms (Figure 1) is applied to the Thornley (1972) model. Internal feedbacks on growth are simulated by making key processes of resource uptake and consumption depend on the internal concentrations of metabolic intermediates which are known to cause such feedbacks. Within this framework, uptake rates, consumption rates and allocation to source and sink tissues are responsive to changes in internal carbon and nitrogen concentrations (Figure 1). The types of feedback mechanisms were chosen to balance each other symmetrically such that, if a feedback is applied to nitrogen, the same type of feedback is implemented to carbon. In order to simplify the system, processes are assumed to be dependent upon local concentrations, for example, carbon uptake rate would be sensitive to changes in leaf carbon and nitrogen but not root carbon and nitrogen. Although there is some experimental evidence for the specific resource status of a compartment and a feedback (e.g. root nitrogen status and leaf growth rate (White et al., 2016)), transport fluxes in the model mean that leaf and root nutrient status are closely coupled.

### 2.1. Developing a framework of feedbacks

The feedbacks (Fig. 1) enable a plant to: increase growth towards sinks when source strength is high for both carbon and nitrogen (Feedbacks 3 & 4); reduce source activity when source strength is high (Feedbacks 1 & 2) and increase carbon source activity when nitrogen source strength is high (Feedback 5) and similarly, for high carbon source strength, increase nitrogen source activity (Feedback 6). By simulating self-shading and root inefficiency with size, the model also has a slow feedback of size on source strength (Eqs. (2) & (3)).

These feedbacks are internal responses which occur with fluctuations in carbon and nitrogen concentration and affect the processes defining growth (carbon uptake; nitrogen uptake; leaf growth; root growth). Each feedback can be implemented mathematically by making the affected process dependent upon the carbon or nitrogen concentration responsible for such a feedback. For instance, feedback 1 alters carbon uptake rate when carbon concentration is high and feedback 5 increases carbon uptake rate with high leaf nitrogen. This means that carbon uptake rate must become dependent upon leaf carbon and leaf nitrogen concentrations. Currently, without any internal feedbacks, carbon uptake rate is assumed to be solely dependent upon atmospheric *CO*_2_ concentration (Eq. (2)). Including the feedbacks, carbon uptake rate would become:

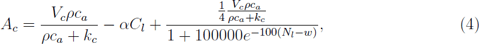

where *αC_l_* is a negative linear feedback on carbon uptake rate when leaf carbon is high (feedback 1), 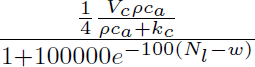is a positive fractional stepwise function of carbon uptake rate when shoot nitrogen is high (feedback 5), *w* = 400*nmolmg^−^*^1^ is the threshold value for shoot nitrogen and root carbon. Nitrogen uptake rate is modelled in the same way, with the same linear function of root nitrogen (feedback 2: high nitrogen reduces nitrogen uptake rate) and with a fractional stepwise function of root carbon (feedback 6: high carbon increases nitrogen uptake rate).

Leaf relative growth rate (*G_l_*) is dependent upon carbon and nitrogen concentration such that:

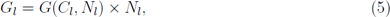

where *G*(*C_l_, N_l_*) is leaf growth rate (see equations 6 & 7) *C_l_*is leaf carbon, *N_l_*is leaf nitrogen. This scales growth rate based on shoot nitrogen to implement a positive feedback on leaf growth rate when nitrogen is high (feedback 3). Similarly, for root growth rate, Eq. (7) is multiplied by root carbon to simulate a positive feedback on root growth when carbon concentration is high (feedback 4).

Since the magnitude of the feedbacks and threshold values aren’t often known, an arbitrary value of 400*nmolmg^−^*^1^ was chosen as a threshold. The supplemental information provides a detailed explanation of how these feedback functions were chosen.

### 2.2. Parameterisation

Parameter values were chosen based on experimental data using a variety of plant species, primarily taken from studies on multiple grasses (Garnier and Laurent, 1994; Tjoelker et al., 2005; He et al., 2006). They are generally relatively fast growing herbaceous species and they would be most appropriate for a model plant species or a crop. This model represents not one particular species but the behaviour of a “generic” plant. Table 1 shows the parameter values used.

**Table 1:** Default parameter values used in modified Thornley (1972) model and definitions.

| Parameter | Value | Units |
| --- | --- | --- |
| $V_c$ - Maximum carbon uptake rate (Sage, 1994) | 40 | $\mu\text{molm}^{-2}\text{s}^{-1}$ |
| $V_n$ - Maximum nitrogen uptake rate (Youngdahl et al., 1982) | 61 | $\mu\text{molkg}^{-1}\text{s}^{-1}$ |
| $k_c$ - Atmospheric $CO_2$ concentration at half of $V_c$ (Farquhar et al., 1980) | 200 | $\mu\text{molmol}^{-1}$ |
| $k_n$ - Soil nitrogen concentration at half of $V_n$ (Youngdahl et al., 1982) | 103 | $\mu\text{M}$ |
| $\rho$ - Ratio of intercellular to atmospheric $CO_2$ concentration (Katul et al., 2000) | 0.7 | $\mu\text{molmol}^{-1}\text{m}^3$ |
| $\beta$ - Ontogenic transport resistance scaling factor (Thornley, 1972) | $V$ | $\text{m}^3$ |
| $r_c$ - Carbon transport resistance | $0.5 \times 10^3$ | $s$ |
| $r_n$ - Nitrogen transport resistance | $1 \times 10^3$ | $s$ |
| $\theta$ - Conversion of plant volume to plant mass (Garnier and Laurent, 1994) | 0.012 | $\text{m}^3(\text{kgmol})^{-1}$ |
| $Y_g$ - Conversion efficiency of substrate to plant material (Kira, 1975) | 0.66 | - |
| $l_0$ - Initial leaf mass | 0.01 | $g$ |
| $r_0$ - Initial root mass | 0.01 | $g$ |
| $C_{l0}$ - Initial leaf carbon concentration of intermediates | 92.8 | $\text{nmolmg}^{-1}$ |
| $C_{r0}$ - Initial root carbon concentration of intermediates | 63 | $\text{nmolmg}^{-1}$ |
| $N_{l0}$ - Initial leaf nitrogen concentration of intermediates | 0.1 | $\text{nmolmg}^{-1}$ |
| $N_{r0}$ - Initial root nitrogen concentration of intermediates | 7.54 | $\text{nmolmg}^{-1}$ |
| $R_1$ - Maintenance leaf respiration (Tjoelker et al., 2005) | 15 | $\text{nmolg}^{-1}\text{s}^{-1}$ |
| $R_2$ - Maintenance root respiration (Tjoelker et al., 2005) | 10 | $\text{nmolg}^{-1}\text{s}^{-1}$ |
| $v$ - Maximum rate of use of carbon and nitrogen for growth | 3600 | $(\text{kgmol})^{-1}\text{m}^3\text{s}^{-1}$ |
| $k_1$ - Michaelis Menten constant for carbon use | 1000 | $\text{kgmolm}^{-3}$ |
| $k_2$ - Michaelis Menten constant for nitrogen use | 1000 | $\text{kgmolm}^{-3}$ |
| $\lambda_1$ - Ratio of nitrogen to carbon atoms within leaf tissue (He et al., 2006) | 0.1 | - |
| $\lambda_2$ - Ratio of nitrogen to carbon atoms within root tissue (He et al., 2006) | 0.2 | - |

Thornley (1972) uses *λ* = 0.11 for the ratio of nitrogen to carbon atoms within plant tissue. In the model here, two separate values are used for leaf (*λ*_1_ = 0.1) and root tissue (*λ*_2_ = 0.2). This reflects the different structural nature of the two tissues (He et al., 2006).

Thornley (1972) simulates plant matter as volume, here, *θ* is used to convert plant volume to plant mass. Thornley (1972) uses *θ* = 0.3*m*^3^(*kgmol*)*^−^*^1^, whilst the new conversion parameter is *θ* = 0.012*m*^3^(*kgmol*)*^−^*^1^ obtained by using a leaf dry density of 10^6^*gm^−^*^3^(Garnier and Laurent (1994)).

The rates of carbon and nitrogen use for plant growth are simulated in the same way to Thornley (1972) but the number of parameter values is reduced by one:

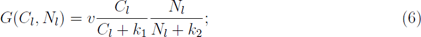

and for root growth:

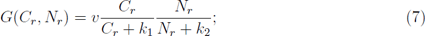

where *v* = *k*, *k*_1_ = 1*/σ_c_* and *k*_2_ = 1*/σ_n_* and assuming that *σ_cn_* = *σ_c_σ_n_* from Eq. (1). These values are parameterised to achieve leaf and root RGR close to 0.3*day^−^*^1^. These values are used throughout this paper (*v* = 3600(*kgmol*)*^−^*^1^*m*^3^*s^−^*^1^ and *k*_1_ = *k*_2_ = 1000*kgmolm^−^*^3^). Although this means that the maximum possible RGR is high (using Eq (S2.11) in supplemental information), with the potential to reach 21.6*d^−^*^1^ (as *C* and *N → ∞*) this value is never reached due to source limitations and is typically less than 0.3*d^−^*^1^.

Additional terms were included in the model to simulate maintenance respiration. Growth respiration is already accounted for in the model by *Y_G_* as the efficiency of growth in converting intermediates into plant tissues but is changed from 0.5 (Thornley, 1972) to 0.66 (dimensionless) (Kira, 1975). The carbon consumed by leaf respiration is subtracted from the intermediate pool of leaf carbon and root respiration from root carbon. Maintenance respiration is simulated as a linear function of plant mass. As plant mass increases, maintenance respiration increases, such that *R*_1_ is leaf respiration (15*nmolg^−^*^1^*s^−^*^1^) and *R*_2_ is root respiration (10*nmolg^−^*^1^*s^−^*^1^) (Tjoelker et al., 2005). All parameter values taken from experimental data were converted to the units used in (Thornley, 1972) and the model output was converted back to the conventionally used SI units for graphical output (see Appendices A and B for table of parameter values and unit conversions).

In order to maximise total plant mass, transport resistance values are reduced from 24 hours to 3 hours. Increasing the transport of resources from sources to sinks reduces the limitations on growth brought about by the concentrations of resources in sink tissues (i.e. root carbon & leaf nitrogen). Similar rates of phloem transport have been measured in **??** and typically vary with herbaceous plant species.

## 3. Results

### 3.1. Atmospheric CO_2_ and nitrogen experiment

By comparing the model output when using two soil nitrogen concentrations and two atmospheric *CO*_2_ concentrations to the patterns observed in experimental data (Coleman et al., 1993; Farage et al., 1998; Rogers et al., 1998; Ainsworth et al., 2003; Butterly et al., 2015), it is possible to determine if the simulated plant with imposed internal feedback mechanisms responds to changes in environment in a similar way to a real plant. Plant growth is simulated for two *CO*_2_ concentrations: 350*ppm* and 700*ppm* for a high soil nitrogen treatment (400*µM*) and a low soil nitrogen treatment (200*µM*).

Figure 3 shows that increasing atmospheric *CO*_2_ concentration leads to a greater total plant mass in both soil nitrogen treatments. Higher soil nitrogen concentrations increases both total plant mass and the effect of *CO*_2_ on total plant mass by 11%. As a consequence, increasing atmospheric *CO*_2_ has a stronger effect on total plant mass with a higher soil nitrogen treatment (high *CO*_2_ creates a 30% change in high nitrogen whilst in low nitrogen the change is 19%). This behaviour replicates a general result from *CO*_2_ and nitrogen experiments (Coleman et al., 1993; Curtis and Wang, 1998; De Graff et al., 2006), such that both nitrogen and *CO*_2_ availability have a positive relationship with plant growth and there is an interaction between *CO*_2_ and nitrogen treatments.

**Figure 3:**
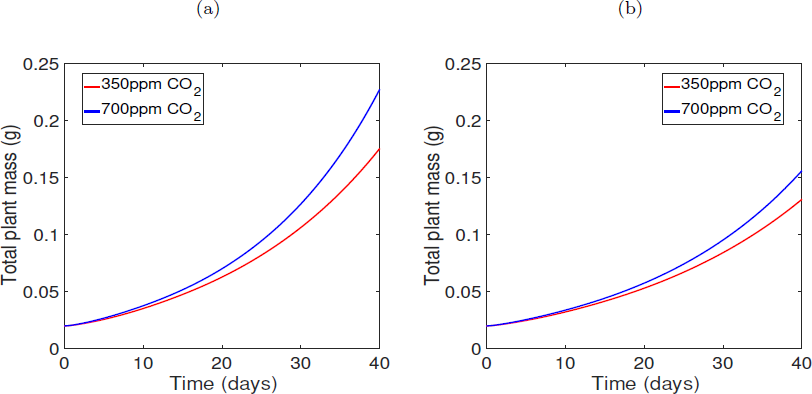
a) The relationship between plant mass over time and two atmospheric *CO*_2_ treatments (350*ppm* and 700*ppm*) with a high soil nitrogen (N=400 *µM*) treatment. b) The relationship between plant mass over time and two atmospheric *CO*_2_ treatments (350*ppm* and 700*ppm*) with a low soil nitrogen treatment (N=200 *µM*). Both ran with initial leaf mass *l*_0_ = 0.01 and root mass *r*_0_ = 0.01 and initial concentrations *C_l_*_0_ = 92.8*nmolmg^−^*^1^*, C_r_*_0_ = 63*nmolmg^−^*^1^*, N_l_*_0_ = 0.1*nmolmg^−^*^1^, *N_r_*_0_ = 7.54*nmolmg^−^*^1^.

Naturally, plants growing in a higher soil nitrogen concentration have a higher percentage of nitrogen in the whole plant (Fig. 4). Increasing atmospheric *CO*_2_ reduces plant nitrogen concentration initially and as the plant continues to grow, a higher atmospheric *CO*_2_ produces a higher percentage of nitrogen than in the ambient control treatment. After 40 days nitrogen concentration is at similar levels for high and low soil nitrogen treatments. When applying a soil nitrogen of 200*µM*, at 40 days, increasing *CO*_2_ raises plant nitrogen percentage slightly from 3.1% to 3.4%. Increasing nitrogen treatment reduces total nitrogen for low (3%) and high (3.4%) *CO*_2_, although this effect is very weak. When plotting percentage of nitrogen within the whole plant against total plant mass (Fig. 4c-d), the same behaviour occurs except that for a low soil nitrogen treatment, the plant produced is smaller. This implies that a higher soil nitrogen treatment produces a larger total plant mass and the weak reduction in nitrogen percentage is a dilution effect arising from a greater plant size.

**Figure 4:**
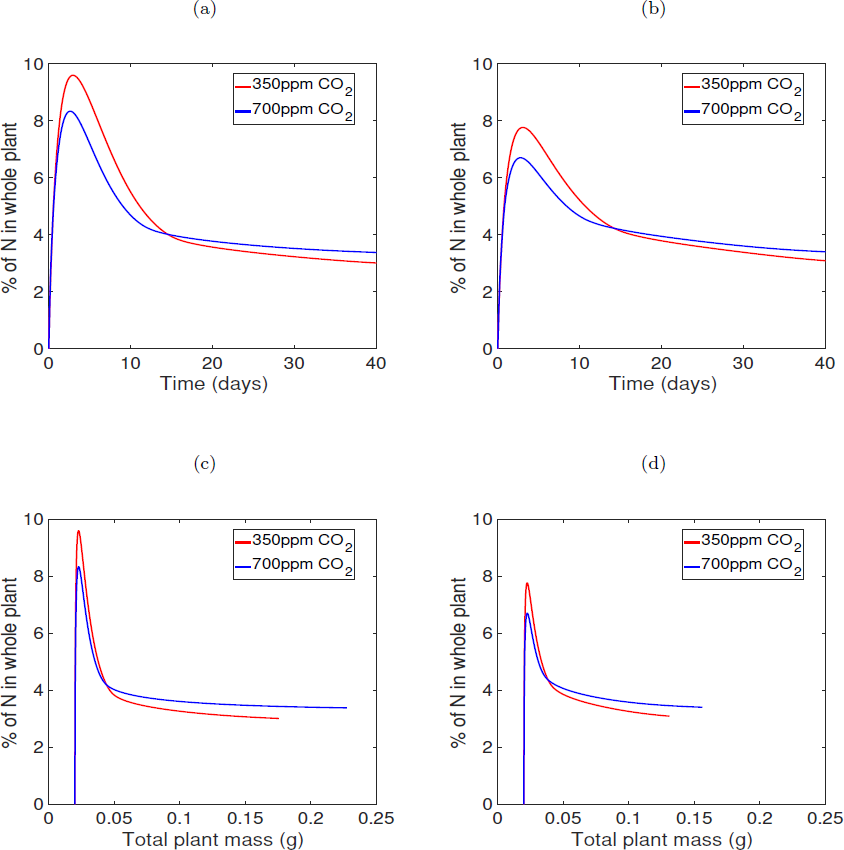
The relationship between nitrogen percentage of total plant mass over 40 days when varying *CO*_2_ treatment (350*ppm* and 700*ppm*) with a) high soil nitrogen (*n* = 400*µM*) b) low soil nitrogen (*n* = 200*µM*). The relationship between nitrogen percentage of total plant mass against plant mass when varying *CO*_2_ treatment (350*ppm* and 700*ppm*) with c) high soil nitrogen (*n* = 400*µM*). d) low soil nitrogen (*n* = 200*µM*). Ran for 40 days with initial leaf mass of 0.01*g* and root mass of 0.01*g* and initial concentrations *C_l_*_0_ = 92.8*nmolmg^−^*^1^*, C_r_*_0_ = 63*nmolmg^−^*^1^*, N_l_*_0_ = 0.1*nmolmg^−^*^1^, *N_r_*_0_ = 7.54*nmolmg^−^*^1^.

Carbon uptake rate depends on atmospheric *CO*_2_, leaf carbon and leaf nitrogen. Given final leaf carbon and nitrogen concentrations when running the model for the combination of high and low *CO*_2_ and soil nitrogen for a period of 40 days, curves of carbon uptake rate over intercellular *CO*_2_ (*A/c_i_*) are produced (Fig. 5a-b). This is done by substituting final leaf carbon and nitrogen concentrations at *t* = 40*days* for both *CO*_2_ and nitrogen treatments into Eq (2) to plot carbon uptake rate against intercellular *CO*_2_ between 0 and 1000*µmolmol^−^*^1^. Creating these plots aids the comparison of responses to environmental change since *A/c_i_* curves are commonly used. Both treatments have a positive effect on carbon uptake rate. High *CO*_2_ levels and soil nitrogen increase carbon and nitrogen uptake. When soil nitrogen is low (200*µM*), atmospheric *CO*_2_ only marginally alters the shape of the *A/c_i_* curve and increasing soil nitrogen maximises the effect of *CO*_2_ on the shape of the *A/c_i_* curve. The effect of atmospheric *CO*_2_ is stronger with a high soil nitrogen treatment for both carbon and nitrogen uptake rate. In the first fifteen days, *CO*_2_ has a large effect on nitrogen uptake rate, as the plant continues to grow, this effect on uptake rate still occurs but diminishes. For a low nitrogen treatment, the effect of *CO*_2_ after 15 days becomes much smaller than when *n* = 400*µM* (Fig. 5c).

**Figure 5:**
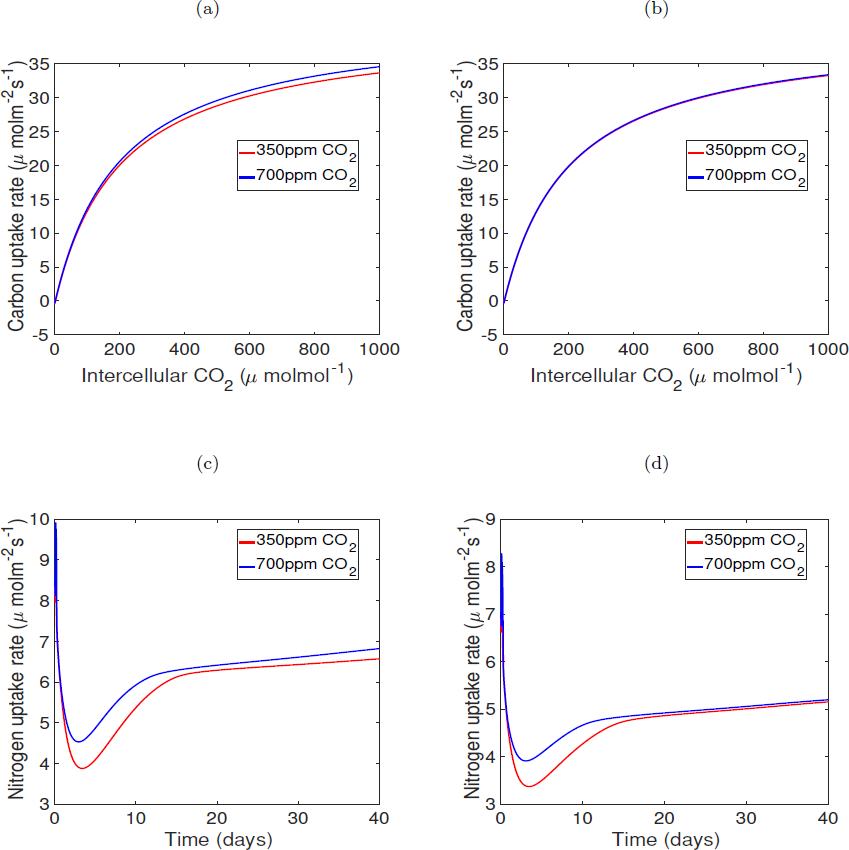
The relationship between carbon uptake rate and intercellular *CO*_2_ for high (700*ppm*) and low (350*ppm*) atmospheric *CO*_2_ a) when soil nitrogen is high (400*µM*). b) when soil nitrogen is low (200*µM*). These curves are created by substituting final leaf carbon and nitrogen concentrations at *t* = 40*days* into Eq (27) to plot carbon uptake rate against intercellular *CO*_2_ between 0 and 1000*nmolmol^−^*^1^. The relationship between nitrogen uptake rate and two atmospheric *CO*_2_ concentrations over 40 days c) when soil nitrogen is high (400*µM*) d) when soil nitrogen is low (200*µM*). The model was run for 40 days with initial leaf mass of 0.01*g* and initial root mass of 0.01*g* and initial concentrations *C_l_*_0_ = 92.8*nmolmg^−^*^1^*, C_r_*_0_ = 63*nmolmg^−^*^1^*, N_l_*_0_ = 0.1*nmolmg^−^*^1^, *N_r_*_0_ = 7.54*nmolmg^−^*^1^.

Figure 6 shows the ratio of root to shoot mass (R:S) as the plant grows for 40 days. The seedlings starts off at equal mass for leaves and roots (*l* = 0.01*g, r* = 0.01*g*). As the plant begins to grow, root growth is favoured but this allocation swaps to leaves very quickly and R:S tends towards 0.3 when soil nitrogen is high and *CO*_2_ is low. Initially for high nitrogen conditions, increasing atmospheric *CO*_2_ has little effect on R:S. High atmospheric *CO*_2_ begins to increase the proportion of roots in relation to leaf mass from 5 days. Low nitrogen treatments increase root growth and the effects of *CO*_2_ treatment emerge sooner than when soil nitrogen is high. Low nitrogen treatment overall produces a higher R:S than a higher nitrogen treatment (Fig. 6b). This reflects the environmental plasticity of the feedback model since when nitrogen availability is high, less roots are produced and when it is lower, more roots are produced. Changes in atmospheric *CO*_2_ have the same effect on R:S under both high and low nitrogen conditions.

**Figure 6:**
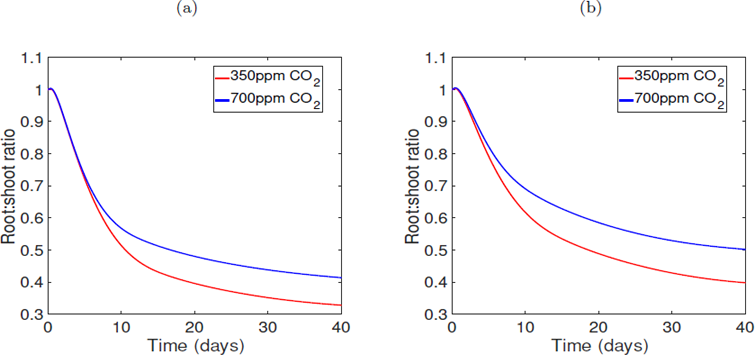
a) The relationship between root:shoot ratio over 40 days and two *CO*_2_ treatments (350*ppm* and 700*ppm*) when soil nitrogen is high (400*µM*). b)The relationship between root:shoot ratio over time and two *CO*_2_ treatments (350*ppm* and 700*ppm*) when soil nitrogen is low (200*µM*). With initial leaf and root mass of 0.01*g* respectively and initial concentrations *C_l_*_0_ = 92.8*nmolmg^−^*^1^*, C_r_*_0_ = 63*nmolmg^−^*^1^*, N_l_*_0_ = 0.1*nmolmg^−^*^1^, *N_r_*_0_ = 7.54*nmolmg^−^*^1^.

The relationships between carbon and nitrogen availability on R:S are a consequence of their relationship with leaf and root relative growth rates. When soil nitrogen is high, increased atmospheric *CO*_2_ simply increases both leaf and root RGR, slightly shifting RGRs. Since the effect of *CO*_2_ is stronger on root RGR than leaf, it leads to an increase in R:S (Fig 6a). When soil nitrogen is low, atmospheric *CO*_2_ also increases individual growth rates but leaf growth is only slightly increased and root growth is increased greatly, rectifying the difference in R:S.

The model was run without any of the internal feedbacks to determine whether it was infact the feedbacks or the (Thornley, 1972) model which is able to respond to changes in environment. The effect of *CO*_2_ and soil nitrogen treatments on total plant mass remains (Fig. 3) when running the experiment on the model without any feedbacks. However, the removal of internal feedbacks allows intermediate nitrogen concentration to increase and therefore nitrogen initially accounts for a much higher proportion of total plant mass than with feedbacks, reaching an unrealistic maximum percentage of 50%. Along with high percentages, increasing *CO*_2_ treatment reduces nitrogen percentage within the plant. Removing the feedbacks produces a similar R:S for all combinations of soil nitrogen and *CO*_2_ treatments such that R:S lies between 0.2 and 0.4. Initially the lower *CO*_2_ treatment has a higher R:S but by 40 days, high *CO*_2_ increases R:S when compared to a lower *CO*_2_ treatment. Due to the lack of internal feedbacks, carbon and nitrogen uptake rate remain constant throughout nutrient availability manipulations (data not shown).

### 3.2. Defoliation and CO_2_ experiment

Defoliation is simulated to investigate how the plant reallocates resources once source size is reduced. To simulate a defoliation experiment, the model was run for 10 days from the same initial conditions used throughout this paper (*l*_0_ = *r*_0_ = 0.01*g*, *C_l_*_0_ = 92.8*nmolmg^−^*^1^*, C_r_*_0_ = 63*nmolmg^−^*^1^*, N_l_*_0_ = 0.1*nmolmg^−^*^1^ and *N_r_*_0_ = 7.54*nmolmg^−^*^1^) with soil nitrogen of 400*µM* . At 10 days, total leaf mass was halved, total internal concentrations and root mass were used as initial conditions and the model was run for another 10 days. This simulation was run for two levels of atmospheric *CO*_2_ (350*ppm* and 700*ppm*).

Defoliation reduces total plant mass but enhances the effect of elevated *CO*_2_ on growth (Fig. 7a). At 20 days, *CO*_2_ increases total plant mass by 12% without any defoliation (Fig. 3a) whereas plant mass is increased by 15% with defoliation (Fig. 7a). Therefore defoliation increases the positive effect of high *CO*_2_ by 3%. This implies that the model with feedbacks is reacting to a halving of the carbon source size (defoliation). Initially, a lower *CO*_2_ treatment produces a plant which is investing more of its resources into leaf growth than high atmospheric *CO*_2_. After defoliation, both treatments appear to be investing into leaf growth at similar rates (Fig. 7b).

**Figure 7:**
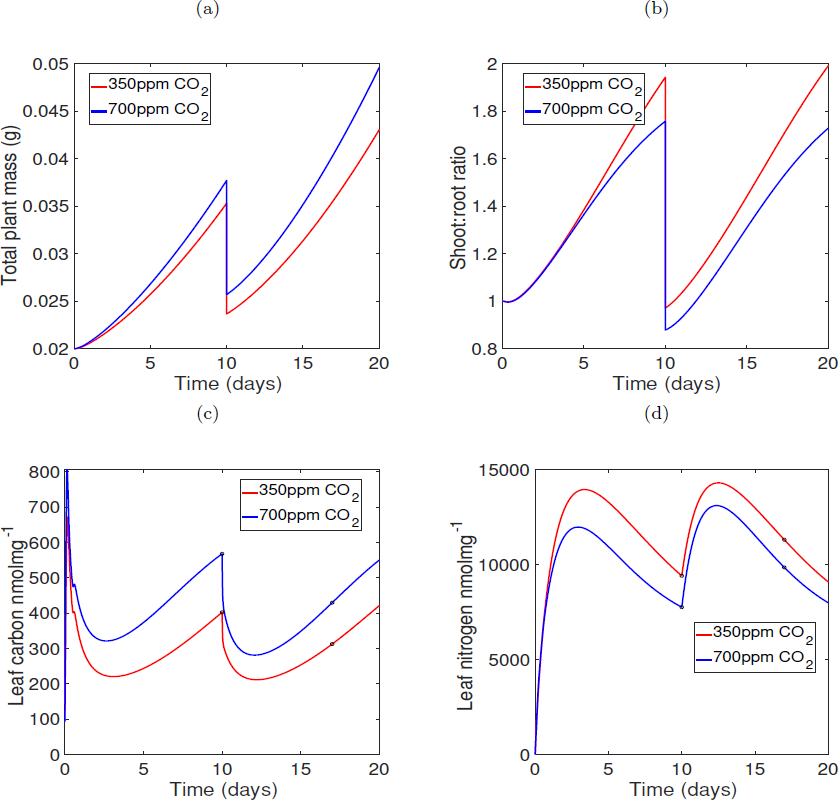
The effect of defoliation (when total leaf mass is halved at day 10) with all internal feedbacks and a high (700*ppm*, blue lines) and low (350*ppm*, red lines) *CO*_2_ treatment on a) Total plant mass over 20 days of growth. b) Proportion of leaf mass compared to root (shoot:root) over 20 days. c) Intermediate leaf carbon concentration for 20 days. d) Intermediate leaf nitrogen over 20 days. Markers signify concentrations of carbon and nitrogen in the leaves at day 10 and day 17 for both carbon and nitrogen plots. All run with soil nitrogen 400*µM* and initial leaf and root mass of 0.01*g* respectively and *C_l_*_0_ = 92.8*nmolmg^−^*^1^*, C_r_*_0_ = 63*nmolmg^−^*^1^*, N_l_*_0_ = 0.1*nmolmg^−^*^1^ and *N_r_*_0_ = 7.54*nmolmg^−^*^1^.

Leaf carbon is reduced and leaf nitrogen increased for both *CO*_2_ treatments, 7 days after defoliation (Fig. 7c-d). For both *CO*_2_ treatments, carbon uptake rate increases initially due to an imbalance between nitrogen and carbon use for growth and respiration and soon reaches a plateau (Fig. 8a). At 7 days after defoliation, carbon uptake rate increases very slightly, with a difference of 0.3% when *CO*_2_ is low and 0.2% when *CO*_2_ is high. Defolia-tion has a much stronger effect on nitrogen uptake rate than for carbon. At 7 days after defoliation, nitrogen uptake rate is reduced by 11.5% when *CO*_2_ is low and 11.7% when *CO*_2_ is high (Fig. 8b).

**Figure 8:**
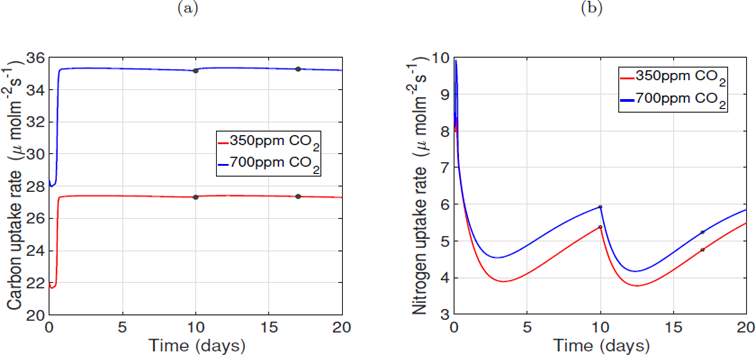
The effect of defoliation (when total leaf mass is halved at day 10) with all internal feedbacks and a high (700*ppm*, blue lines) and low (350*ppm*, red lines) *CO*_2_ treatment on a) Carbon uptake rate over 20 days of growth. b) Nitrogen uptake rate over 20 days of growth. Markers signify carbon and nitrogen uptake rate at day 10 and day 17. All ran with soil nitrogen 400*µM* and initial leaf and root mass of 0.01*g* respectively and *C_l_*_0_ = 92.8*nmolmg^−^*^1^*, C_r_*_0_ = 63*nmolmg^−^*^1^*, N_l_*_0_ = 0.1*nmolmg^−^*^1^ and *N_r_*_0_ = 7.54*nmolmg^−^*^1^.

Carbon concentration decreases 7 days after defoliation when simulating the experiment without any internal feedbacks on growth and uptake. This also applies for intermediate nitrogen concentration such that after 7 days, nitrogen is higher. Removing the feedbacks on intermediate concentration therefore does not alter the effect of defoliation on concentration. However, without feedbacks defoliation does not alter carbon and nitrogen uptake rates since in the absence of feedbacks they are only dependent upon external *CO*_2_ and nitrogen respectively.

## 4. Discussion & conclusions

The aim of this paper was to determine whether a framework model of feedback mechanisms sensitive to changes in carbon and nitrogen responds to changes in the environment in a similar way to experimental data (Coleman et al., 1993; Farage et al., 1998; Rogers et al., 1998; Ainsworth et al., 2003; Butterly et al., 2015) and to highlight areas of uncertainty where more empirical data is needed. This was carried out by simulating two levels of atmospheric *CO*_2_ and two soil nitrogen treatments in factorial combination and a defoliation experiment where the total above ground biomass was halved after 10 days and growth was simulated for an additional 10 days. These simulations showed that the model qualitatively reproduces the *CO*_2_ and nitrogen interaction often observed experimentally, whereby the positive effect of atmospheric *CO*_2_ on growth is magnified with high nitrogen and diminished by low nitrogen (Demmers-Derks et al., 1998; Coleman et al., 1993). The model produces higher nitrogen concentrations and an increase in net carbon uptake rate against intercellular *CO*_2_ with elevated atmospheric *CO*_2_, which does not match experimental results. Additionally the model reproduces the experimental finding that defoliation increases the positive effect of high *CO*_2_ on growth (Ryle and Powell, 1992; Wand and Midgley, 2004). Moreover, increased atmospheric *CO*_2_ leads to a larger simulated root:shoot ratio and increased soil nitrogen leads to a lower proportion of roots than either ambient *CO*_2_ and nitrogen treatments respectively. Imposing defoliation in the model produces higher leaf nitrogen, lower leaf carbon and higher carbon uptake rates as observed experimentally (Von Caemmerer and Farquhar, 1984; Rogers et al., 1998; Eyles et al., 2013). Therefore, a model which simulates internal feedbacks on source and sink strengths with changes in carbon and nitrogen is able to reflect key behaviours observed in experiments that manipulate source strength. This model takes us one step closer towards modelling the mechanisms responsible for allocation.

The model accurately shows that elevated soil nitrogen increases the positive effect of in-creased *CO*_2_ on total plant mass (Demmers-Derks et al., 1998; Coleman et al., 1993). This arises because low soil nitrogen imposes a limit on how much biomass can be produced (i.e. sink limitation), however this does not feed back onto photosynthetic rate (Fig. 5a-b). The model simulates a slight raising of the *A/c_i_* curve in elevated compared with ambient CO2 in high N, contradicting experimental observations (Coleman et al., 1993; Ainsworth et al., 2003). However, this is something that is theoretically predicted at high CO2 (Sage, 1994; Woodrow, 1994). The increased availability of *CO*_2_ increases nitrogen uptake rate (via feedback 5) and consequently nitrogen availability. Since carbon and nitrogen concentrations drive leaf and root relative growth rate, the immediately available concentrations of carbon and nitrogen are used for new tissue production. Therefore the other feedback mechanisms have acted in place of feedback 1 (high carbon reduces carbon uptake rate) and the intermediate carbon pool has not been able to accumulate high enough to trigger a reduction in carbon uptake rate.

The model simulations in this paper show that high levels of atmospheric *CO*_2_ promote an increase in root:shoot ratio, whilst increasing soil nitrogen reduces root:shoot ratio. This is to be expected within the model as high levels of carbon increase root growth and conversely high levels of nitrogen increase leaf growth via the internal feedback mechanisms. Individual studies have shown increases (Lacointe, 2000) and decreases (Butterly et al., 2015) in root:shoot ratio with elevated *CO*_2_. However, a meta-analyis by Ainsworth et al. (2002) found no change in root:shoot ratio. Butterly et al. (2015) also found that increasing nitrogen availability increases root:shoot ratio whilst Vicente et al. (2015) and Dybzinski et al. (2011) found that increasing nitrogen reduces root:shoot ratio. Therefore empirical evidence conflicts on whether root:shoot ratios should increase or decrease from changes in *CO*_2_ and soil nitrogen availability. An emergent property of the model is the decline in root:shoot ratio with age and is seen commonly in plants (Poorter et al., 1988; Negrini et al., 2020). Another emergent property of the model is that low N availability produces small plants with reasonable intermediate concentrations of carbon and nitrogen (Wu et al., 2007) rather than the alternative of producing larger plants with a diluted nitrogen content and consequences of functioning (Vos et al., 2005). Reducing the ratio of nitrogen to carbon atoms within plant tissue (*λ*) in the model should allow this alternative growth strategy to occur.

Defoliation enhances the positive effect of high atmospheric *CO*_2_ within the model. Removing half of the total leaf mass after 10 days makes the plant source limited, reducing carbon source strength whilst nitrogen source strength remains the same. This leads to an increase in leaf nitrogen concentration due to higher availability and a reduction in leaf carbon. The high levels of leaf nitrogen triggers the feedbacks of high N to increase carbon uptake rate whilst reducing nitrogen uptake rate and an increase in leaf growth rate. These responses entirely match experimental observations (Ryle and Powell, 1992; Wand and Midgley, 2004; Von Caemmerer and Farquhar, 1984; Rogers et al., 1998; Eyles et al., 2013). Furthermore, the framework model is able to respond to defoliation in a robust way.

A number of assumptions were made in order to simulate source-sink interactions, highlighting areas of focus for further experimental testing. Firstly, the rate of tissue growth is assumed to depend on carbon and nitrogen concentration in the model however, this may not necessarily be the case. Additional data is required on the rate of carbon and nitrogen use for growing plant tissue and how this changes over a growing period. Secondly, the form of each feedback function used in the model were determined mathematically based on (Anonymous, 2019). However the way in which carbon or nitrogen concentrations trigger feedbacks on growth processes is largely unknown. Most experiments do not focus on the time it takes for these feedbacks to affect plant growth or the threshold values or sensitivities of signal and response to pool size, they simply address the time scale of their measurements. There is a clear lack of knowledge on the time it takes from when concentrations are sensed, to gene expression for the induction of enzymes for a reaction, to the change in shoot:root ratio.

Nunes et al. (2013) show that when imposing sink limitation by reducing temperature or low nitrogen and supplying arabidopsis with sucrose, T6P increases which increases the use of carbon for growth. T6P inhibits the expression of SnRK1 in order to increase growth processes. They present a starvation threshold for sucrose of 3*µmolg^−^*^1^ (fresh weight) and T6P of 0.3 *−* 0.5*nmol*T6P*g^−^*^1^ (fresh weight). This paper determines a threshold value that sugar must surpass in order for the rate of use of carbon for growth to be promoted. To better understand these mechanisms, not only are threshold values are needed to be obtained experimentally but also the rates at which these feedbacks occur. It is unclear whether these threshold values represent a triggered switch like behaviour or whether the response is more gradual and continuous. Furthermore, the strength of the feedback responses on growth processes are also unknown and are simulated to be dependent upon carbon or nitrogen concentration within the model.

This model can be extended to include other known feedback mechanisms for instance, sugars have been shown to negatively influence the loading of sugars, altering the rate of transport of substrate between leaves and roots (Chiou and Bush, 1998; Vaughn et al., 2002; **?**). The feedbacks chosen here only simulate responses to growth and uptake rates with high levels of carbon and nitrogen, whereas other known behaviours are in response to low concentrations. For instance, when sugars are scarce meristem growth stops (Lastdrager et al., 2014). SnRK1 protein kinase is present when sugars are low and this is responsible for suppressing growth (Baena-Gonáalez et al., 2007; Polge and Thomas, 2007; Halford and Hey, 2009; Baena-Gonáalez, 2010; Ghillebert et al., 2011) but sucrose can also stimulate SnRK1 (Baena-Gonáalez, 2010). Low sugars can also stop the transcription of nitrate reductase (Stitt and Krapp, 1999; Klein et al., 2000; Kaiser et al., 2002; Reda, 2015). Additionally, leaf nitrogen concentration sets a limit to the maximum capacity of carbon assimilation through a relationship with carboxylation rate (**?**). This presents other feed-backs which could be incorporated in the the model. Further detailed analysis is needed on which feedbacks are more important than others, for instance, what is the smallest number of feedbacks required to simulate reasonable responses to changes in environment? Are additional feedbacks required to make the model behave more reasonably?

Overall, the results of this paper show that the model with internal feedback mechanisms based on internal carbon and nitrogen concentrations is able to reproduce most of the behaviours seen in experiments varying carbon and nitrogen availability qualitatively. It reproduces more of the observed behaviours than a model without feedbacks and is able to regulate internal concentrations of carbon and nitrogen. The framework presented here extends the work of other models which have simulated dependency of source or sink activity on carbon and nitrogen (Bartelink, 1998; Hunt et al., 1998; Dunbabin et al., 2002; ÅAgren et al., 2012; Pao et al., 2018; Shaw and Cheung, 2018) by including more types of feedbacks which alter both source and sink activity. By incorporating more of the observed feedbacks, this work provides a closer representation of allocation processes. This model provides a tool to investigate the dynamics between internal feedback mechanisms which control the allocation of biomass and uptake of external resources, and could be incorporated into crop models to give process-based mechanistic feedbacks between carbon and nitrogen resource availability. Furthermore, this paper sharpens questions about the physiological mechanisms underpinning resource allocation and highlights new research directions. Understanding the mechanisms behind allocation can provide new areas of focus to manipulate the net primary productivity of plants.

## Supporting information

Suppmental information

