## Supplementary material for "Implementing a framework of carbon and nitrogen feedback responses into a plant resource allocation model": Suppmental information

### Methods S1

Five functions with varying levels of complexity representing each feedback type were compared by using a dimensionless four compartment model of leaf mass, root mass and intermediate carbon and nitrogen (Anonymous, 2019). The dimensionless analysis enables easy comparison of how to model internal feedbacks of carbon or nitrogen concentration on growth processes. The function which best represents each feedback under three environmental conditions (equal maximum carbon and nitrogen uptake rates, maximum carbon uptake rate is lower than nitrogen and maximum nitrogen uptake rate is lower than carbon) is used. These were chosen based on the following constraints:

- The function successfully produces the behaviour of the feedback;
- RGR tends towards zero as time increases and is always positive;
- Growth rate tends towards a constant steady state value, therefore representing linear growth;
- RGR is within an order of magnitude compared to maximum RGR without feedbacks;
- Carbon and nitrogen uptake rates are within an order of magnitude compared to their maximum rates.

It was determined that the best function to simulate negative feedbacks on carbon and nitrogen uptake rates when carbon and nitrogen concentrations are high is a linear one for three environmental cases (equal uptake rates, low carbon uptake and low nitrogen uptake). A scalar function best describes the behaviour of a positive feedback onto leaf and root growth rates when nitrogen and carbon is high. For a positive feedback on uptake rates, a fractional stepwise function best simulates a positive response with high concentrations for all three environmental conditions. This is how each feedback is modelled in this paper.

**Methods S2**

Total leaf carbon concentration depends on the uptake rate of carbon via photosynthesis, the export of carbon to the roots and the cost of producing new plant material. The Thornley model is thus:

$$\frac{d(V_lC_l)}{dt} = A_cV_l - \frac{\beta(C_l - C_r)}{r_c} - V_lF(C_l, N_l); \quad (1)$$

$$\frac{d(V_lN_l)}{dt} = \frac{\beta(N_r - N_l)}{r_n} - V_l\lambda F(C_l, N_l); \quad (2)$$

$$\frac{d(V_rC_r)}{dt} = \frac{\beta(C_l - C_r)}{r_c} - V_rF(C_r, N_r); \quad (3)$$

$$\frac{d(V_rN_r)}{dt} = A_nV_r - \frac{\beta(N_r - N_l)}{r_n} - V_r\lambda F(C_r, N_r); \quad (4)$$

$$\frac{dV_l}{dt} = \theta Y_gV_lF(C_l, N_l); \quad (5)$$

$$\frac{dV_r}{dt} = \theta Y_gV_rF(C_r, N_r); \quad (6)$$

where $V_l$ and $V_r$, are shoot and root volume, respectively ($m^3$), $C_l$ and $N_l$ are carbon and nitrogen concentration per unit leaf volume ($kgmolm^{-3}$), $C_r$ and $N_r$ are carbon and nitrogen concentration per unit root volume, respectively ($kgmolm^{-3}$), $t$ is time ($s$), $A_c$ and $A_n$ are carbon and nitrogen uptake rate per unit leaf and root volume respectively ($kgmolm^{-3}s^{-1}$), $\beta$ is a parameter that describes how transport scales with plant size ($m^3$), $r_c$ and $r_n$ are carbon and nitrogen transport resistance ($s$), $\lambda$ is the N:C ratio of atoms in the plant (dimensionless), $\theta$ is the conversion of dry matter to volume ($m^3(kgmol)^{-1}$), $Y_g$ is the efficiency of converting carbon and nitrogen concentrations into plant material (dimensionless) and $F(C_l, N_l)$ and $F(C_r, N_r)$ are functions for the rate of use of substrate (i.e. sink activities $kgmols^{-1}$). In this current form, Equations (3) - (6) represent the total amount of leaf carbon and nitrogen and root carbon and nitrogen within the whole plant.

$C_lV_l$ represents the amount of carbon in all above ground biomass and therefore $C_l$ is the amount of carbon per $kg$ of leaf (i.e. the concentration of carbon intermediates available for growth). For simplicity, the product rule is applied to the differentials in Equations (3) - (6), to obtain size-independent substrate concentration equations. The model becomes:

$$\frac{dC_l}{dt} = A_c - \frac{\beta(C_l - C_r)}{r_c V_l} - (1 + C_l \theta Y_g) F(C_l, N_l); \tag{7}$$

$$\frac{dN_l}{dt} = \frac{\beta(N_r - N_l)}{r_n V_l} - (\lambda + N_l \theta Y_g) F(C_l, N_l); \tag{8}$$

$$\frac{dC_r}{dt} = \frac{\beta(C_l - C_r)}{r_c V_r} - (1 + C_r \theta Y_g) F(C_r, N_r); \tag{9}$$

$$\frac{dN_r}{dt} = A_n - \frac{\beta(N_r - N_l)}{r_n V_r} - (\lambda + N_r \theta Y_g) F(C_r, N_r); \tag{10}$$

$$\frac{dV_s}{dt} = \theta Y_g V_l F(C_l, N_l); \tag{11}$$

$$\frac{dV_r}{dt} = \theta Y_g V_r F(C_r, N_r). \tag{12}$$

Overall plant growth is defined by Equations (B.11) and (B.12). This implies that growth is dependent upon a function for the use of carbon and nitrogen per volume, the conversion of dry matter to volume ($\theta$), and a plant mass conversion efficiency ($Y_g$), which is the equivalent to growth respiration. $F(C, N)$ is a function representing the rate of use of substrate for leaf and root growth (Eq. (B.13)). The amount of nitrogen used for growth of new tissue, $F(C, N)$ is multiplied by the N:C ratio of atoms in plant tissue ($\lambda$). The solutions to the model (Equations ( B.7) - (B.12)) are found using Ode23s in MATLAB which is a single step solver for stiff systems by calculating the Jacobian matrix of the system at each time step.

$F(C, N)$ represents sink activity and depends on substrate limitation. This is an increasing function of both carbon and nitrogen. The maximum growth rate represents sink capacity (when C and N $\rightarrow \infty$), which is given by $k/\sigma_{cn}$. For constant N, $N = N_0$, $F(C, N_0)$ is a Michaelis-Menten function of C.

$$F(C, N_0) = \frac{k N_0 C}{1 + \sigma_n N_0 + (\sigma_c + \sigma_{cn}) C}. \tag{13}$$

As $C \to \infty$,

$$F(C, N_0) \to \frac{kN_0}{\sigma_c + \sigma_{cn}}. \tag{14}$$

The rate of change of substrate concentration is dependent upon its import (via uptake if a source, via transport if a sink), the transport to other compartments, and the use of substrate for growth.

The transport term within the model enables a slow feedback on the tissue concentration pools. The transport term is divided by shoot / root volume. In order for dimensions to balance within the model, it is essential for the ontogenic scaling factor for transport to be equivalent to total plant volume ($\beta = V$, where $V = V_l + V_r$). This enables transport resistance to scale with plant volume such that the rate of transport is slower in larger plants. For leaf carbon, the transport term from the leaf to the root is:

$$\text{Carbon transport rate} = \frac{V_l + V_r}{V_s} \times \frac{(C_l - C_r)}{r_c}. \tag{15}$$

This can be written as

$$\text{Carbon transport rate} = \frac{(C_l - C_r)}{r_c} + \frac{V_r}{V_l}\frac{(C_l - C_r)}{r_c}, \tag{16}$$

illustrating the slow feedback on concentrations. The first term in Equation (B.8) is a general transport term that moves the difference in concentration from the carbon source to the sink. The second term depends on the ratio of root:shoot volume. Hence, if there is much higher root volume than leaf volume, an additional amount at the value of the size difference is transported to the leaves. This is an internal feedback within the model to counteract any imbalances in the root:shoot (R:S) ratio.

**Methods S3**

Increasing maximum nitrogen uptake rate increases total plant mass after 20 days of growth (Fig. 4a). As $V_n$ increases towards 1000 $\mu mol kg^{-1} s^{-1}$

total plant mass begins to plateau. This is because nitrogen and carbon are required for growth, as nitrogen uptake tends towards 1000 $\mu molkg^{-1}s^{-1}$, carbon uptake rate becomes limiting. Maximum carbon uptake rate also has a positive relationship with total plant mass and starts to plateau as nitrogen uptake becomes limiting (as $V_c$ tends towards 1000 $\mu molm^{-2}s^{-1}$). The rate at which total dry plant mass reaches a plateau when varying maximum carbon uptake is not as defined as for nitrogen. There are a lot of parameter values which determine the ratio of use of carbon and nitrogen, setting the ratio of nitrogen to carbon atoms within leaf and root tissue to be more equal ($\lambda_1 = \lambda_2$) produces a smoother curve. Although the relationship between carbon uptake rate and total plant mass is similar to nitrogen, carbon uptake rate produces roughly twice the total plant mass when compared to varying total nitrogen uptake rate. This can be explained by the parameterisation of the model. A much larger proportion of plant mass is made up of carbon for leaves and roots than nitrogen (there is ten-times more carbon than nitrogen in leaves ($\lambda_1 = 0.1$) and five-times more carbon than nitrogen in the roots ($\lambda_2 = 0.2$)).

The model results are very sensitive to initial seedling size (which would arise ecologically through variation in seed size (Fig. 4c)). Total dry plant mass after 20 days is proportional to initial seedling mass, a relationship that arises from exponential growth. The range of parameter values for leaf and root maintenance respiration is limited as maintenance respiration has a negative relationship with total plant mass and leaf and root RGR. When maintenance respiration is low, the plant is able to carry out exponential growth. As respiration increases, the amount of available carbon for tissue production is limited until the amount of carbon needed for maintenance becomes larger than the amount imported via photosynthesis. At this point RGR becomes negative. This represents an unrealistic parameter space since plants would alter internal processes to avoid a negative RGR. For the parameters used (see appendix), maintenance respiration for leaf and root tissue cannot exceed $56nmolg^{-1}s^{-1}$ equally (Fig. 4d). The ratio between carbon uptake rate and maintenance respiration must be large enough to sustain growth.

Transport resistance has a negative relationship with total plant mass, such that increasing resistance reduces total plant mass (Fig not shown). This effect doesn't occur when transport resistance is almost non-existent ($0.01s$), where increasing transport resistance increases total plant mass. This is a consequence of the transport resistance parameter, since it divides the amount of resource being transported. Therefore really small values cause instability within the model. Once transport resistance increases past $1s$, total plant mass decreases. Varying parameter values responsible for the rate of substrate use for growth (RGR) has little effect on total plant mass since RGR is limited by the availability of carbon and nitrogen intermediates given the default parameter set. This implies that in its current state (without feedbacks), the plant is source limited. Increasing the ratio of nitrogen to carbon atoms that make up leaf and root mass decreases total plant mass. Additionally, increasing the parameters which represent the concentration of carbon and nitrogen, when carbon and nitrogen uptake rate is half of its maximum respectively ($k_c$ and $k_n$), reduces total plant mass because doing so reduces RGR for given levels of carbon and nitrogen.

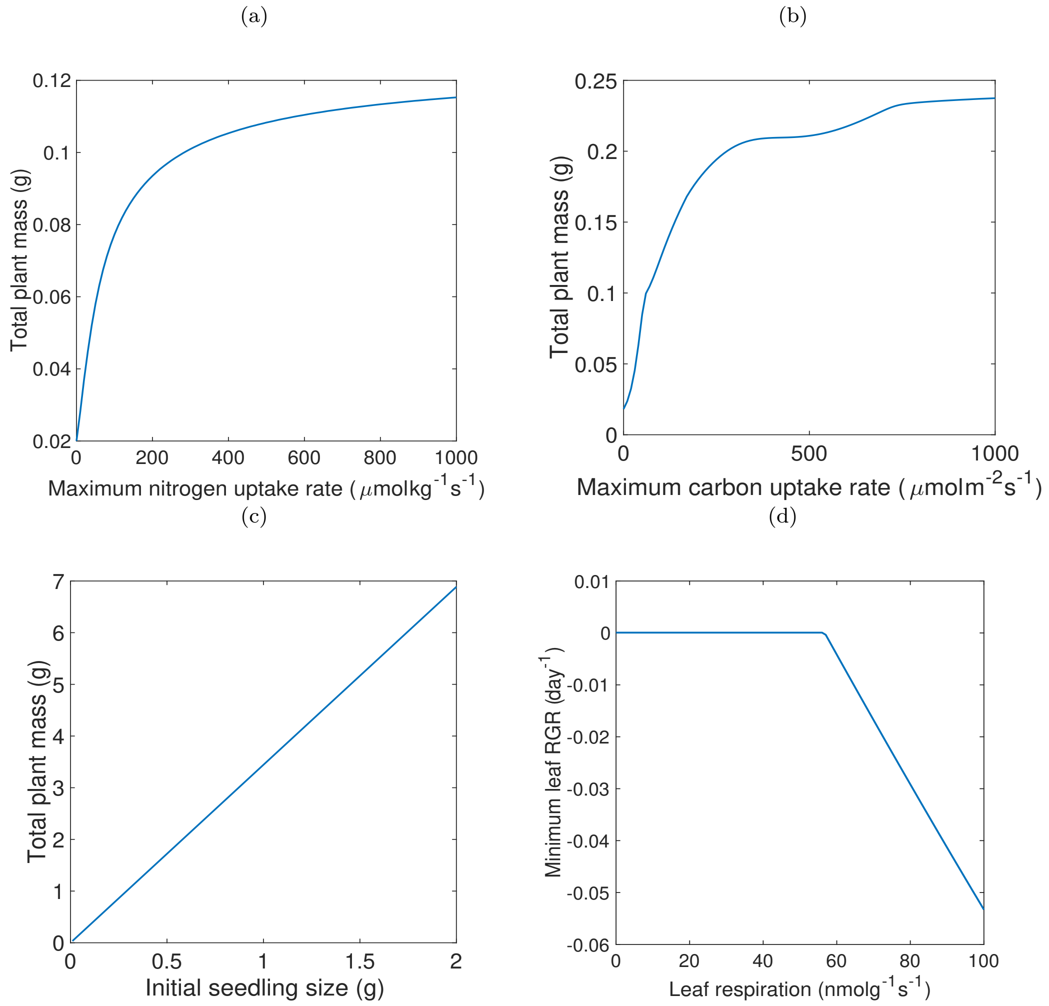


Figure 1: a) The relationship between maximum nitrogen uptake rate and total plant mass after 20 days of growth. b) The relationship between maximum carbon uptake rate and total plant mass. c) The relationship between initial seedling size (initial leaf mass and initial root mass) and total plant dry mass after 20 days of growth. d) The relationship between leaf maintenance respiration and minimum leaf RGR. Leaf respiration and RGR is equal to root respiration and RGR for this subfigure. All figures produced using default parameter values (see appendix) with initial leaf and root mass of $0.01g$ respectively grown for 20 and initial concentrations $C_{l0} = 92.8nmolmg^{-1}, C_{r0} = 63nmolmg^{-1}, N_{l0} = 0.1nmolmg^{-1}$, $N_{r0} = 7.54nmolmg^{-1}$.
